# Socio-affective touch expression database

**DOI:** 10.1101/161729

**Authors:** Haemy Lee Masson, Hans Op de Beeck

## Abstract

Socio-affective touch communication conveys a vast amount of information about emotions and intentions in social contexts. In spite of the complexity of the socio-affective touch expressions we use daily, previous studies addressed only a few aspects of social touch mainly focusing on hedonics, such as stroking, leaving a wide range of social touch behaviour unexplored. To overcome this limit, we present the Socio-Affective Touch Expression Database (SATED), which includes a large range of dynamic interpersonal socio-affective touch events varying in valence and arousal. The original database contained 26 different social touch expressions each performed by three actor pairs. To validate each touch expression, we conducted two behavioural experiments investigating perceived naturalness and affective values. Based on the rated naturalness, 13 socio-affective touch expressions along with 12 corresponding non-social touch events were selected as a complete set, achieving 75 video clips in total. Moreover, we quantified motion energy for each touch expression to investigate its intrinsic correlations with perceived affective values and its similarity among actor-and action-pairs. As a result, the touch expression database is not only systematically defined and well-controlled, but also spontaneous and natural, while eliciting clear affective responses. This database will allow a fine-grained investigation of complex interpersonal socio-affective touch in the realm of social psychology and neuroscience along with potential application areas in affective computing and neighbouring fields.

## Introduction

Humans frequently express emotions and intentions through nonverbal communication (1). Among these nonverbal communication behaviours, facial expression received the most attention, yielding more than a hundred different facial expression databases (e.g., GEMEP Corpus^1^ and MMI Facial Expression Database^2^). Yet, interpersonal touch also conveys a vast amount of information such as socio-affective state and physical and psychological closeness between the interacting people (2), to which humans are exposed even before birth (3). Thus, an expression through touch is a potent and hard-to-ignore means of conveying a message, notwithstanding the individual and cultural variation in terms of how, how often, and with whom, touch communication is employed. Moreover, interpersonal touch often has a strong emotional valence. Touch as positive affect plays an important role in making the receiver feel support, reassurance, affection or sexual attraction (4). In contrast, touch as negative affect can convey anger, frustration or disappointment (5). Hence, interpersonal touch often is affective touch (2, 6). Over the past decade, psychologists and neuroscientists have increasingly studied the role of touch in social context, how the emotions were conveyed through touch, and its neural basis. Despite of the importance and high frequency of touch communication and increasing interest in this topic, however, stimuli created for these studies poorly represent the actual phenomenon of socio-affective touch, and are mostly limited to video clips showing simple events such as a hand being stroked, slapped or brushed (7-14). These studies therefore only address a very small part of "affective touch" since humans use much more complex and varied interactions to express affective intentions (both positive and negative) by touch. A wider range of more complex interpersonal affective touch events has so far only been addressed in studies using static images (for example, using pictures of kissing, hugging, and hitting as stimuli: (15-18). However, there has been no attempt to implement complex dynamic affective touch stimuli in this domain. Crucially, none of the aforementioned studies varies their stimuli for levels of different valence and arousal, leaving important evaluative dimensions open (19).

To allow for more rigorous study of the social and affective impact of touch, we developed the Socio-Affective Touch Expression Database (SATED), with the aim to cover a larger range of dynamic interpersonal socio-affective touch actions, which span two primary affective dimensions, valence and arousal. To reach this general aim, we started by designing a systematically defined, well-controlled socio-affective touch space. Next, we validated the affective values and characterised the physical aspects of these stimuli using a behavioural experiment and a computer vision algorithm respectively. Lastly, we created additional video clips containing situations in which objects were touched without inducing specific emotions (neutral touch towards objects) while the movements used for object-based touch (non-social) itself were matched with the movements used in interpersonal socio-affective touch.

## Materials and Methods

### Creation of a stimulus set of socio-affective and non-social touch videos

The major goal of the current study is to create and validate dynamic interpersonal socio-affective touch video clips that span a large range of touch communication events, as well as their corresponding object-based non-social touch events. In order to cover two primary affective dimensions, valence and arousal, we included negative, neutral, and positive touch situations, which are associated with low (calm) to high arousal (exciting). Examples of negative situations include punching and pushing a person on his/her arm. Examples of positive situations include different forms of gently stroking a person (for the purpose of consolation or flirting) and hugging. Examples of neutral situations include shaking hands as a greetings and nudging the arm to get someone’s attention.

Six actors (three actor-pairs) without any professional acting experience volunteered in recordings. The idea behind recruiting non-professional actors was to capture rather spontaneous and life-like touch expressions rather than trained prototypical acting. Each actor pair consisted of one male and one female actor who were familiar to each other, and between whom at least neutral touch expressions had been frequently used in a real life situation. Therefore, actors did not feel awkward to use touch communication during recordings. On the day of recordings, actors wore either black or grey long-sleeves shirts so as not to induce biased reactions towards the clothing styles between actors while also providing a high visual contrast for distinguishing the actors from each other and the background. All the videos were recorded in the same place where an empty white wall was used as a background. The camera used for recordings has resolution of 1280 (width) × 720(height) pixels with frame rate 29 frames per second. The camera was mounted to a tripod at a distance of 1.71 metre from the actors.

Each pair of actors performed 26 different socio-affective touch scenes in front of the camera. A method-acting protocol was used in which actors were instructed to read a scenario description provided beforehand and to think about a similar situation in their life in order to call up the emotions and actions that one needs in the filming situation (20). Here, we asked actors to use a specific action (hug, stroke, hit or shake) and duration (approximately three seconds) for each touch expression. In this way, we could still control the action and duration while making other factors such as speed and pressure of the touch as natural as possible.

Table 1 lists the scenarios describing the situations that evoke intended touch expressions in the final stimulus set. This list only contains a basic set of stimuli that were retained after the validation test (See results) and includes both socio-affective and corresponding object-based touch events. It contains 13 socio-affective events x; 3 actor-pairs, and in addition 12 object-based touch events. Full scenarios of the original larger set of socio-affective touch events (26 events × 3 actor-pairs) can be found in S1 Table.

**Table 1.** Scenarios of the final set of touch events.

| Video Number | Scenario | Valence | Action | Stimulus Name |
| --- | --- | --- | --- | --- |
| 1, 14, 27 | You are in the airport. You and your partner have not seen each other for 6 month. You hug each other as soon as you see each other. | Positive | Hug | Hug1_p |
| **2, 15, 28** | Your partner looks somehow lovelier today. You want to give a hug to express how much you love this person. | Positive | Hug | Hug2_p |
| **3, 16, 29** | Your sibling just ended the long-term relationship. You want to console him (her) by hugging. | Positive | Hug | Hug3_p |
| **4, 17, 30** | You want to flirt with him (her) by stroking his (her) arm with intimacy. | Positive | Stroke | Str1_p |
| **5, 18, 31** | Your friend is crying. You want to console this person by stroking on his (her) arms. | Positive | Stroke | Str2_p |
| **6, 19, 32** | You hold hands of your partner with affection. | Positive | Hold | Hold1_p |
| **7, 20, 33** | You want to get an attention from your colleague (who could not hear you calling) by tapping his (her) shoulder. | Neutral | Tap | Tap1_neu |
| **8, 21, 34** | You just found out that your partner cheated on you. You are very disappointed and angry. You ask him (her) how and why this happened aggressively while shaking his (her) body. | Negative | Shake | Sha1_n |
| **9, 22, 35** | Your sibling committed crime. You are sad and angry since you cannot understand why he (she) made such a horrible decision. You firmly ask him (her) how and why this has happened while shaking his (her) arm. | Negative | Shake | Sha2_n |
| **10,23,36** | Your partner said that you two are over from now on. He or she is about to walk away and never see you again. You cannot let this person go. You cannot let this person go. You grip this person's arm desperately. | Negative | Grip | Gri1_n |
| **11,24,37** | You are in the metro. You need to get off from the metro. There is somebody blocking you. You nudge your way by slightly removing somebody's arm blocking you. | Negative | Nudge with elbow | Nudg1_n |
| **12,25,38** | Your sibling tries to be silly by making a weird gesture. You are annoyed. You want to stop him (her) by poking in the ribs. | Negative | Nudge with elbow | Nudg2_n |
| **13,26,39** | Your sibling always make the same mistakes. You slap his (her) arm to make him (her) realize that you are annoyed and that you expect something better. | Negative | Slap | Slap1_n |
| **40,52,64** | Carry the water bottle. | Neutral | Hug | Hug2_ob |
| **41,53,65** | Carry the big box. | Neutral | Hug | Hug3_ob |
| **42,54,66** | Examine the texture. | Neutral | Stroke | Str1_ob |
| **43,55,67** | Remove possible wrinkles from the suit jacket hung on the hanger. | Neutral | Stroke | Str2_ob |
| **44,56,68** | Carry the basket. | Neutral | Hold | Hold1_ob |
| **45,57,69** | Ring the desk bell to get attention. | Neutral | Tap | Tap1_ob |
| **46,58,70** | Empty the dust from the plastic box. | Neutral | Shake | Sha1_ob |
| **47,59,71** | Shake the cocktail in the mixer. | Neutral | Shake | Sha2_ob |
| **48,60,72** | Shake the Champaign bottle before opening it. | Neutral | Shake | Sha3_ob |
| **49,61,73** | Pass by a door while elbowing it closed. | Neutral | Nudge | Nudg1_ob |
| **50,62,74** | Push with arm to close the door. | Neutral | Nudge | Nudg2_ob |
| **51,63,75** | Remove dust from the carpet. | Neutral | Slap | Slap1_ob |

For the object-based touch events that cover the non-social touch category, we included situations where similar motions were required to interact with a particular object. For example, the touch scene in which the fabric of clothing is stroked was included as a motion match to the touch event where one actor strokes the other’s arm. It should be taken into account that we created object-based touch scenarios after the validation test on social stimuli to match the action between social and non-social video clips. The idea behind having object-based touch videos as a set of control stimuli is that such a contrast condition might be useful for various experimental applications, such as neuroimaging and neuropsychological experiments. Therefore, actors returned back to perform the object-based touch events on a later date. They were instructed to wear the same clothes that they used for the social-affective touch events. Care was taken that the object-based touch scenarios would not induce distinctive emotional responses, hence expected to be evaluated as “relatively neutral” in terms of valence ratings.

For all touch events, actors were allowed to practice the event beforehand until it felt natural to perform. Once they agreed to start, they performed each scene after hearing the voice of the director who recorded the video clips saying, “start”. During recordings, actors were allowed to produce facial and verbal expressions if necessary in order to keep the naturalness. Both audio and face information were discarded during the recording and the editing, with the focus being only on the movement of torso. All actors were able to perform all touch actions in spite of different level of difficulty between stimulus events (e.g. neutral vs. aggressive expressions). Fig 1 shows representative still frames from the selected dynamic touch expressions.

**Fig 1.**
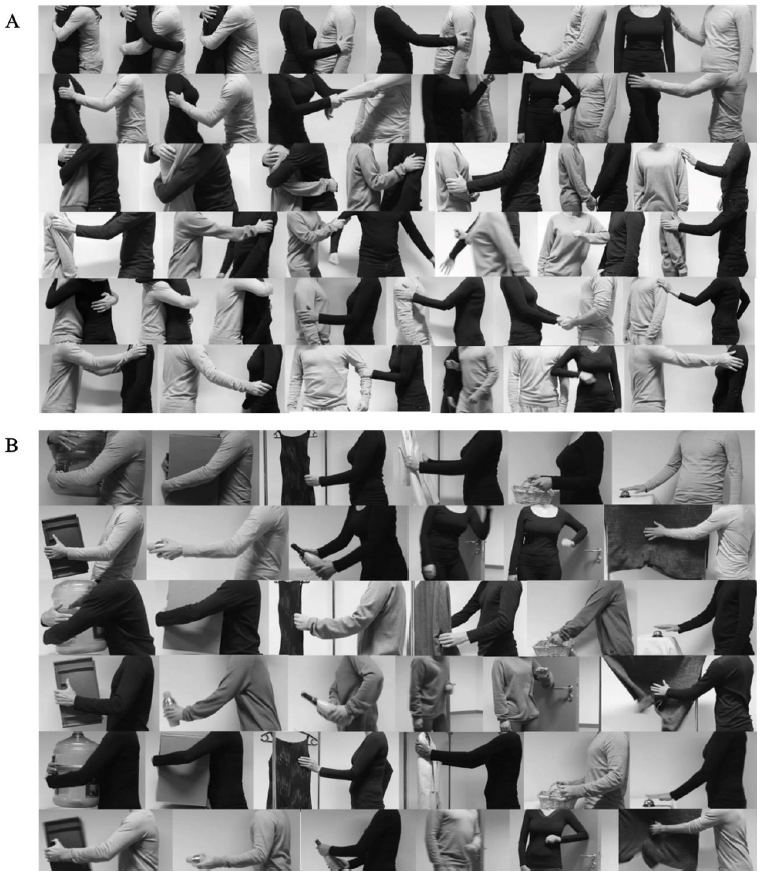
Socio-affective and non-social touch stimuli. A) The figure shows representative still frames from the basic set of socio-affective stimuli, showing different types of interpersonal touch events (positive (the first six stimuli in the 1st, the 3rd and the 5th rows), neutral (the last stimuli in the 1st, the 3rd and the 5th rows) and negative events (six stimuli in the 2nd, the 4th and the 6th rows)). B) The figure shows representative still frames from the matched object-based non-social stimuli, showing different interactions with various objects. The touch expressions are available as dynamic video clips.

### Validation of stimulus set

The purpose of this stimulus validation experiment was to test if the affective aspects of the social video clips are perceived as positive or negative as intended, while also measuring their naturalness so as to exclude artificially performed socio-affective touch scenes. To achieve these aims, we recruited participants who judged valence, arousal and naturalness of each stimulus (Experiment 1). After the selection of social touch stimuli was made, object-based touch stimuli were created and validated along with the social ones on valence and arousal ratings (Experiment 2). Affective responses of object-based touch events were examined to test whether object-based touch events could serve the function as control stimuli. Lastly, the association between motion energy and affective characteristics were investigated.

### Participants

First, 11 right-handed adults (6 female, mean age 27.2) rated the social video clips based on valence, arousal, and naturalness (Experiment 1). After selecting the optimized stimuli set through the validation test, 22 right-handed adults (10 female, mean age 26) were recruited to rate the valence and arousal for the final set of stimuli including both social and non-social video clips (Experiment 2). Neither previous neurological nor psychological histories were reported among participants. Written informed consent was provided before the experiment. The study was approved by the ethical review board of KU Leuven (G-2016 06 569).

### Stimuli and experimental design

In Experiment 1, the stimuli consist of video clips displaying interpersonal socio-affective touch scenes in the context of pleasant to unpleasant situations were validated during the experiment. The social stimuli included 26 scenes with positive (N=10), neutral (N=6) and negative (N=10) touch interactions from three sets of actors (78 stimuli in total). Participants watched the videos, containing socio-affective touch events and evaluated them by rating its valence (“How pleasant is the touch?”), arousal (“How (emotionally) arousing is the touch?”) and naturalness (“How natural is the video clip?”) of the touch scene on a 7-point Likert-like scale using the keyboard. This validation experiment would lead us to exclude stimuli with low naturalness ratings and to be able to evaluate if expected valence ratings were met across the social stimuli.

In Experiment 2, object-based touch stimuli were created and validated together with the final set of 13 selected social scenarios. As mentioned above, the object-based touch stimuli were to function as a set of control stimuli that matched the actions of the social stimuli. Similar to the validation experiment described above, participants watched the videos, containing both socio-affective touch and object-based touch events, and evaluated them by rating its valence and arousal on a 9-point Likert-like scale using Self-Assessment Manikin (SAM) (21).

In both experiments, each video was shown twice in a random order on a 23-inch LCD monitor (at a resolution of 640 (width) × 360 (height)) located at 0.5 metres from the participant. The experiment was controlled by Psychophysics Toolbox Version 3.0.12 (PTB-3) (22) in MATLAB. Fig 2 illustrates the procedures of the two experiments.

**Fig 2.**
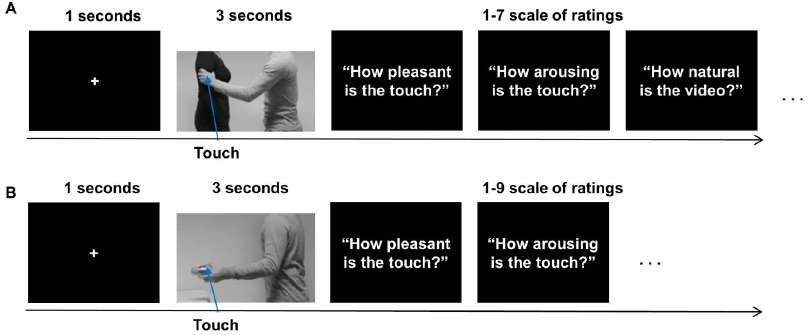
Experimental design. (A) The figure illustrates the experimental design for the validation test in Experiment 1. Participants were asked to answer three questions [valence, arousal and naturalness] using 1-7 scale after watching social touch videos for 3 seconds. (B) The figure shows how the object-based videos were validated along with the social videos in Experiment 2. Only valence and arousal were asked using 1-9 scale with SAM.

### Analysis: Naturalness

First, the average group naturalness ratings were calculated across stimuli (26 stimuli × 3 actor-pairs). We measured median naturalness of each stimulus across three actor-pairs to exclude any stimulus perceived artificially by participants. Our criteria for the exclusion were any stimuli exhibit either median rating lower than scale 4 (neutral) or are not significantly higher than scale 3 (slightly artificial) with a student t-test.

### Analysis: Intra-, inter-subject and inter-actor consistencies

Using correlational analysis with Spearman rank-order correlations, we measured the intra-observer (across repetitions within participants), inter-observer (across participants), and inter-stimulus (across performances of three actor-pairs) consistencies.

### Analysis: Affective responses

A correlational analysis between valence and arousal was performed to confirm whether our data exhibited a U-shaped distribution as previously reported (21). Moreover, we evaluated if positive stimuli such as hugging were rated pleasantly and negative videos such as slapping unpleasantly. Importantly, we measured if object-based touch events induced distinctive emotional responses despite they were intended to be “neutral” in terms of valence ratings.

### Analysis: Low level visual motion features

Physical parameters such as the amount and type of motion are intrinsically connected to the motion displayed in the videos. For example, slapping involves short transients of high-speed motion. Thus, identifying motion intensity enables us to understand its possible links with action and emotional responses. We identified the magnitude of the motion energies to characterize its interaction with affective properties of touch video clips. Moreover, we examined consistency across actor-pairs in terms of consumed total motion energies to identify the similarities among the actor-pairs while performing the touch expression in a quantitative manner. Lastly, we compared the motion intensity of socio-affective touch with those of object-based touch. An Optical flow algorithm based on the Lucas-Kanade method was used that determine the flow field between two consecutive frames of dynamic stimuli under several constraints (23, 24). With the algorithms, we obtained magnitude of motion energy by summing the absolute values of estimated velocities across the frames, which implies how much motion energy was consumed to perform particular actions (e.g. stroking or shaking) regardless of direction of the motion (Fig 3).

**Fig 3.**
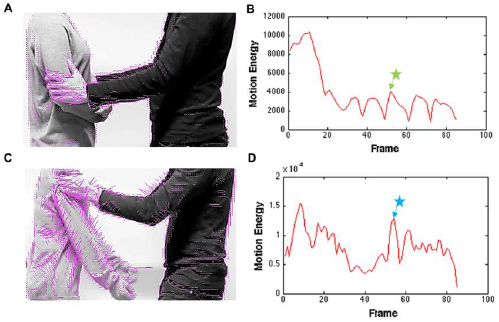
Motion energy. (A) and (C) An example frame from the video sequences of positive (A) and negative (C) touch event with inserted pink lines. The length and the orientation of pink lines indicate speed and direction of the pixel movements respectively. (B) and (D) Total motion energy of all pixels from the first frame to the last frame. The star signs with arrows in (B) and (D) indicate which frames are shown in (A) and (C) respectively.

## Results

### Validation experiment

**[H3] Experiment 1: Naturalness of the stimuli**

The results of the naturalness judgments indicated that the video clips were perceived naturally overall (mean rated naturalness 4.86 on a 7-point Likert-like scale; 1-very artificial, 4-neutral, 7-very natural). Fig 4 demonstrates the naturalness ratings of each of the 26 scenarios, averaged across the three actor-pairs. Based on the results, we excluded the stimuli rated rather artificially from the set of socio-affective stimuli and from the further analyses. Hence, stimulus number 16 (airport check) and 25 (forcefully hugging) were subject to be excluded since the median naturalness of those videos was below threshold (See the red line in the Fig 4, see S1 Table for description of the stimuli). Moreover, stimulus number 8 (shaking hands with joy for celebration), 21 (an aggressive punch), 22 (an aggressive push) are excluded after the student's t-test since the naturalness of those stimuli was not significantly above the scale 3 (See the yellow dashed line in the Fig 4). Most of the excluded stimuli include negative scenarios, which reflects the finding that it was easier for the actors to perform positive scenes (mean 5.19) than negative ones (mean 4.36) (t(10)=3.47, p<0.01). After removing the stimuli mentioned above, average naturalness judgments became 5.03, implying that the 21 remaining scenarios resulted in videos that were perceived naturally.

**Fig 4.**
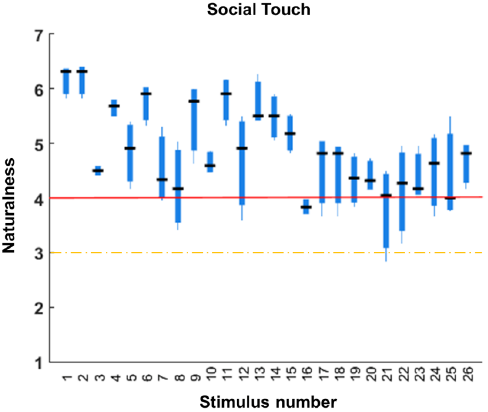
Group naturalness ratings. The figure demonstrates the group naturalness ratings of each stimulus acted by three actor-pairs. The central black lines indicate the median naturalness ratings. The top and bottom edges of the boxes illustrate the 75th and 25th percentiles respectively while the whiskers show the range of the naturalness ratings. The red line and orange dashed line indicate the naturalness ratings 4 (neutrally perceived) and 3 (slightly artificially perceived) respectively.

**[H3] Experiment 1: Intra-, inter-subject and inter-actor consistencies**

The results indicated that participants were consistent about their ratings between two repetitions (valence median RHO=0.95, range 0.98∼0.89; arousal median RHO=0.81, range 0.93∼0.46; naturalness median RHO=0.74, range 0.87∼0.29) Based on the critical values of the Spearman`s ranked correlation coefficient table (25), a correlation coefficient above 0.435 is considered to be significant at the alpha level 0.05 for N = 21, where N is the number of correlated scenarios (Fig 5A).

**Fig 5.**
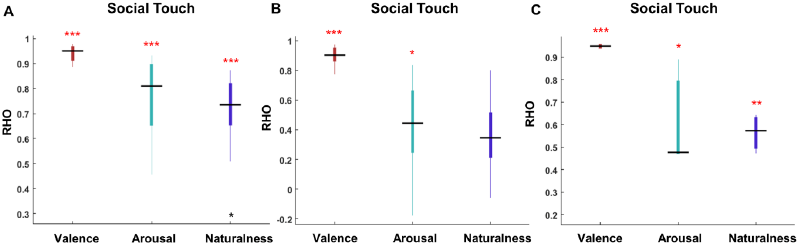
Intra-, inter-subject and inter-actor consistencies. The figure illustrates the intra-subject variability (A), the inter-subject variability (B) and the inter-actor variability (C) on valence (dark red), arousal (emerald) and naturalness (purple) ratings. In all figures, the central black lines indicate the medians of Spearman correlations. The top and bottom edges of the boxes illustrate the 75th and 25th percentiles respectively while the whiskers show the range of the rank correlation coefficients. The black asterisk indicates the outlier in (A). The single red asterisk indicates the statistical significance at P<0.05, two red asterisks denotes statistical significance at P<0.01, and three red asterisks show statistical significance at P < 0.001.

We also found high correlation on valence ratings between participants, suggesting high inter-observer consistency (median RHO=0.9, range 0.97∼0.78). In case of arousal (median RHO=0.44, range 0.84∼-0.18), we found mild correlation across participants. However, based on Zar’s statistical inferences (25), the correlation of rated naturalness (median RHO=0.35, range 0.8∼-0.06) across participants was not significant, implying individual variability on perceived naturalness (Fig 5B).

Lastly, we measured the inter-actor consistency to validate whether the same scenarios performed by three different pairs of actors yielded the similar affective responses and naturalness ratings. Again, we found high correlation on valence (median RHO=0.95, range 0.95∼0.94) and naturalness (median RHO=0.57, range 0.64∼0.47) ratings across stimuli sets performed by three different pairs of actors, suggesting high inter-actor consistency. In the case of arousal (median RHO=0.48, range 0.89∼0.48), we found mild correlation across actor-pairs. Based on Zar’s statistical inferences (25), we could infer that participants perceived each scenario performed by three different actor-pairs rather similarly in all ratings (Fig 5C).

**[H3] Experiment 1: Affective responses of the stimuli**

We investigated the affective responses to the 21 scenarios with a sufficiently high naturalness rating. Importantly, we were able to observe the typical U-shape relationship between arousal and valence (21) obtained from quadratic function fitting of the averaged group ratings (Fig 6A). All the positive touch events were perceived pleasantly (mean rating 5.79; 1-very unpleasant, 4-neutral, 7-very pleasant) whereas the negative touch events were rated unpleasantly (mean rating 2.48; 1-very unpleasant, 4-neutral, 7-very pleasant). Fig 6B presents the group valence ratings across 21 touch events (See S1 Table for description of the stimuli).

**Fig 6.**
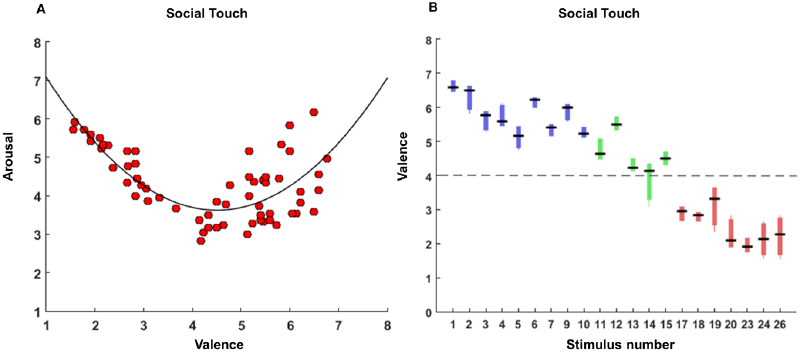
Affective responses of social stimuli. (A) The scatter plot and fitted quadratic function between group valence and arousal ratings of 63 stimuli [21 touch events × 3 actor-pairs] present the typical U-shape distribution. The black curve indicates the fitted polynomial curve of degree 2 using a least-squares regression between valence and arousal. (B) The boxplots show group valence ratings of 21 touch events performed by three actor-pairs. The central black lines indicate the medians of group valence ratings. The top and bottom edges of the boxes illustrate the 75th and 25th percentiles respectively while the whiskers show the range of the valence ratings. Different colours indicate the expected valence of each touch event [blue for the pleasant, green for the neutral and red for the unpleasant events]. Black dash line is dividing the stimuli between the pleasant and unpleasant touch events at the point of rating 4 (neutral valence), indicating the stimuli above the line are perceivably pleasantly while stimuli below the line are perceive unpleasantly.

### Selection of a basic set of stimuli and object-based touch

Based on the results described above, we selected 6 positive, 1 neutral and 6 negative touch scenes as a basic set of stimuli (See Table 1, Fig 1). We did not select most of the neutral stimuli since they were rated rather more pleasantly than neutrally—for example, see in Fig 6B the stimuli 11 (shake hands for greeting), 12 (hugging for greeting) and 15 (waking somebody up).

After the selection of the basic set of stimuli, we created the object-based non-social touch videos that had matched hand gestures with social videos such as hugging, stroking, and shaking. We recorded a total of 36 stimuli [12 motion matched scene × 3 actor pairs]. It should be noted that one of the social videos where both actor are running to hug (See Table 1, stimulus name “Hug1_p”) could not be matched since an object cannot run and hug a person. Thus, a basic set of stimuli consists of 13 social and 12 object-based scenarios, performed by different actors.

**[H3] Experiment 2: Affective responses to the basic set of social and object-based video clips**

We measured valence and arousal of each selected social touch video and its corresponding object-based touch video. Similar to the results described above (See Fig 6B), positive touch events were perceived as pleasant (mean rating 7.38; 1-extremely unpleasant, 5-neutral, 9-extremely pleasant) while negative events were perceived as unpleasant (mean rating 2.74; 1-extremely unpleasant, 5-neutral, 9-extremly pleasant). Importantly, object-based touch events were rated as neutral (mean rating 4.87; 1-extremely unpleasant, 5-neutral, 9-extremly pleasant). Fig 7 shows both group valence and arousal ratings of the 25 stimuli [13 social touch and 12 corresponding object-based touch], averaged across actors. Student's t test revealed that stimulus number 3 (Str1_ob; t(21)=2.63, p<0.05), 7 (Sha1_ob; t(21)=-3.3, p<0.01), 10 (Nudg1_ob; t(21)=-6.18, p<0.001) and 11 (Nudg2_ob; t(21)=-5.41, p<0.001) were significantly different from the neutral value 5 (See red asterisks in Fig 7A). Yet, all stimuli were rated significantly (p<0.05) differently from the mean ratings of positive touch (7.38) and negative touch (2.74), showing that object-based touch events were *relatively* neutral in terms of valence ratings. We also compared the rated arousal between social and non-social touch. The results revealed that social videos (except neutral touch, stimulus name “Tap1_neu” see Fig 7B and Table 1) were rated more arousing (mean rating 6.11; 1-extremely calm to 9-extremely exciting) than non-social videos (mean rating 2.5) (t(11)=13.15,p<0.001), as we intended.

**Fig 7.**
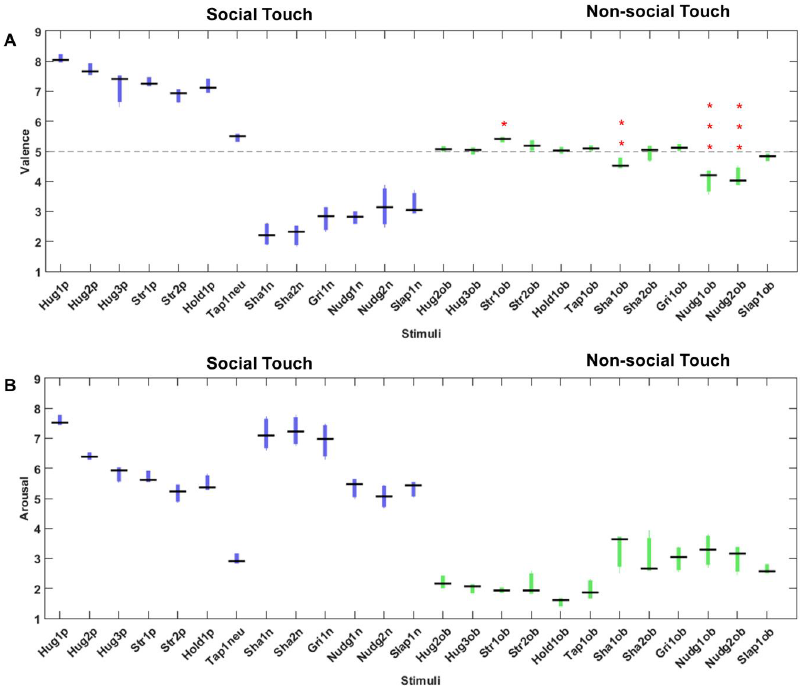
Affective responses of social and non-social stimuli. The boxplots show group valence (A) and arousal ratings (B) of 25 touch events performed by three actor-pairs for the social videos and 6 actors for the non-social videos. The central black lines indicate the medians of ratings. The top and bottom edges of the boxes illustrate the 75th and 25th percentiles respectively while the whiskers show the range of the ratings. Different colours indicate the distinction between social and non-social touch stimuli (blue for the social touch and green for the non-social touch events). Black dash line in the (A) is dividing the stimuli between the pleasant and unpleasant touch events at the point of rating 5 (neutral valence), indicating the stimuli above the line are perceived pleasantly while stimuli below the line are perceived unpleasantly. The single red asterisk indicates the statistical significance at P<0.05, two red asterisks denotes statistical significance at P<0.01, and three red asterisks show statistical significance at P < 0.001. X-axis indicates the name of stimuli (See Table 1).

### Motion energy and its association with affective responses

First, rank correlational analysis on motion energy across actor-pairs revealed that similar motion energies characterized the three pairs of actors across a social video set (13 videos per pair of actors) (median RHO=0.74, range: 0.81∼0.69) and an object-based video set (12 videos) (median RHO=0.62, range: 0.67∼0.46). Moreover, the results showed that motion energy between social and non-social videos across actor-pairs are highly associated (RHO=0.64, range: 0.85∼0.61) as we intended since the motions were matched across the stimuli. Based on the study of Zar (25), a correlation coefficient above 0.56 is considered to be significant at the alpha level 0.05 with an N of 13 (number of video clips which we correlated, 0.587 with an N of 12 for the non-social videos). In all cases, similar patterns of motion energy were used among actors during recordings across scenario (hugging across actor-pairs) and matched body motion (hugging somebody and hugging the box).

Secondly, we used student’s two-sample t test between social videos (N=39) and non-social videos (N=36) to evaluate the differences on motion energy. The results revealed that motion energy was not significantly different between social and non-social videos (t(73)=-0.1, p=0.92).

Lastly, correlational analysis between motion energy and affective ratings of all stimuli (N=75) showed a significant, expected association between the rated arousal of touch events and their physical motion energy (RHO=0.55, p<0.001). Moreover, rated valence and motion energy also showed a weak association (RHO=-0.31, p<0.01). The scatter plots in the Fig 8 illustrate the association between motion energy and affective ratings.

**Fig 8.**
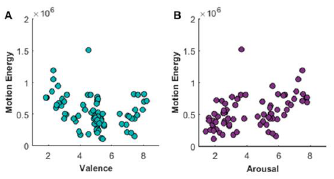
Motion energy and its association with affective responses. The scatter plots illustrate weak negative association between rated valence and motion energy (A) and moderate positive association between rated arousal and motion energy (B). Both x-axis represent 9-point Likert-like scale of ratings.

## Discussion

The aim of this study was to devise a reliable touch expression database that covers a large repertoire of interpersonal socio-affective touch. To achieve this goal, we created and validated a systematically defined, well-controlled socio-affective touch space, including negative, neutral, and positive situations with both low (calm) to high arousal (exciting) content. As control stimuli we also added video clips containing situations in which objects were touched without inducing specific emotions while matching each non-social action to a social touch event.

The results from the validation experiments enabled us 1) to eliminate artificially performed interpersonal touch scenes, 2) to measure whether interpersonal touch communication exhibits clear and replicable affective values and 3) to define intrinsic physical characteristics so as to control for or at least understand potentially unavoidable low-level physical parameters.

In spite of a large degree of individual variances in rank order according to naturalness of the social touch expressions, the majority of video clips (21 scenarios out of 26), performed by non-professional actors basing on written scenarios, were perceived naturally. Although we did not provide either an utterly artificial or a highly natural touch expression as references before starting with the naturalness judgements task, based on the results that describe our participants being consistent across repetitions, we conjecture that participant consistently relied on their own criteria for the judgements without varying them during the task, providing us with a useful exclusion criterion for the artificially performed touch scenes. Each touch expression performed by three different actor-pairs was also rated similarly on its naturalness, proving a shared level of difficulty depending on the given scenario among actor-pairs. Indeed, informal discussions with the actors suggested that the negative scenarios were more difficult to perform, yet they were perceived naturally. Given the results on rated naturalness, there is some ground to claim that the final socio-affective touch database (after the exclusion of some of the original videos) is composed of spontaneous performed, natural and life-like interpersonal touch expressions.

We also tested inter-and intra-observer consistencies on valence and arousal ratings. The results illustrated that participants were consistent on their ratings across the repetitions while conforming to others, showing both intra and inter-observer agreements on affective judgements. Moreover, rated affective values on each video clip performed by three different actor-pairs were also similar, indicating that our scenarios, used during recordings, successfully anchored the actor-pairs to perform in a way that positive touch events were perceived as pleasant, whereas negative touch events were perceived as unpleasant. When it comes to arousal, more variability within and across participants was found as compare to valence, which conforms with the findings of a previous study that investigated valence and arousal with stimuli from the frequently used International Affective Picture System (26). Importantly, the typical U-shape relationship between arousal and valence was observed for our stimulus set (21), proving the validity and reliability of rated arousal with respect to valence. Overall, the results demonstrate that our socio-affective database elicit clear and replicable affective responses which span primary affective dimensions (19).

After selecting the basic set of stimuli, we created and validated corresponding object-based non-social touch stimuli. Here, our aim was to create object-based touch stimuli that did not evoke specific emotion during observation. Indeed, our findings illustrate that object-based touch events are relatively neutral in terms of pleasantness as compare to social touch events. In the case of arousal, social touch turned out to be more arousing than object-based non-social touch, showing that interpersonal touch is emotionally more intense.

Lastly, we systematically measured motion energy across frames to characterize intrinsic physical properties of a particular action accompanied during social and non-social touch situation. Our findings revealed that the same video clips (e.g., hugging someone) acted by three different actor-pairs required similar amount of motion energy, indicating inter-actor consistency on performance across given scenarios. Moreover, the same actions (e.g., hugging someone vs. hugging a box) were similarly characterized by this motion parameter regardless of socio-affective components embedded in the social touch situations. In a similar vein, total amount of motion energy consumed between social and non-social videos does not differ, implying inter-action consistency on consumed motion energy.

We also observed a positive association between rated arousal and motion energy in line with the expectation based on previous findings that demonstrated accompanied high motion energy in emotionally charging movie scenes displaying anger, horror and excitement, implying an intrinsic link between the two (27, 28).

Although our database, SATED, carries an unprecedented large range of dynamic interpersonal touch communication, we should admit that it still leaves out other parts of affective touch events. For example, real-life touch events regularly evolve a face (e.g., stroking or slapping a face). However, revealing the facial identity of actors could modulate the affective judgements on touch communication due to face attractiveness. Based on a previous study, attractiveness of a touch communicator can influence on perceived pleasantness of affective touch (29). Thus, in this database, we controlled for this variable by limiting the touch information to the torso region without any face information, and by all actors wearing similar type of clothes (See Methods).

In summary, we created a large set of interpersonal touch expression database, the Socio-Affective Touch Expression Database (SATED), containing both social and non-social events. This validated database can facilitate research on human touch communication, serving as an advanced set of stimuli, ultimately enabling us to properly investigate more complex interpersonal touch communication. We envision that our database can be used in various domains such as affective computing and social neuroscience.

Our database, SATED, is freely available in Open Science Framework (https://osf.io/8j74m/) for scientific use.

## Author contributions

H.L.M. and H.O.d.B. conceived the project and designed the experiments. H.L.M. conducted experiments and analysed data. H.L.M. and H.O.d.B. wrote the paper.

## Acknowledgements

We gratefully acknowledge the acting performance of anonymous volunteers and the help of Nicky Daniels for assistance with recordings.

## Footnotes

1 Banziger T, Scherer KR. Introducing the geneva multimodal emotion portrayal (gemep) corpus. Blueprint for affective computing: A sourcebook. 2010 Sep 23:271-94.

2 Valstar M, Pantic M. Induced disgust, happiness and surprise: an addition to the mmi facial expression database. InProc. 3rd Intern. Workshop on EMOTION (satellite of LREC): Corpora for Research on Emotion and Affect 2010 May 21 (p. 65).

